# NSUN6, an RNA methyltransferase of 5-mC controls glioblastoma response to Temozolomide (TMZ) via NELFB and RPS6KB2 interaction

**DOI:** 10.1101/2021.08.11.455957

**Authors:** Chidiebere U Awah, Jan Winter, Claudiane M Mazdoom, Olorunseun O. Ogunwobi

## Abstract

Nop2/Sun RNA methyltransferase (NSUN6) is an RNA 5 - methyl cytosine (5mC) transferase with little information known of its function in cancer and response to cancer therapy. Here, we show that NSUN6 methylates both large and small RNA in glioblastoma and controls glioblastoma response to temozolomide with or without influence of the MGMT promoter status, with high NSUN6 expression conferring survival benefit to glioblastoma patients and in other cancers. Mechanistically, our results show that NSUN6 controls response to TMZ therapy via 5mC mediated regulation of NELFB and RPS6BK2. Taken together, we present evidence that show that NSUN6 mediated 5mC deposition regulates transcriptional pause (by accumulation of NELFB and the general transcription factor complexes (POLR2A, TBP, TFIIA, TFIIE) on the preinitiation complex at TATA binding site to control translation machinery in glioblastoma response to alkylating agents. Our findings open a new frontier into controlling of transcriptional regulation by RNA methyltransferase and 5mC.

## Introduction

RNA methylation is the new frontier in understanding of epitranscriptomics, which represents a new layer in the control of genetic information. The messenger RNA (mRNA), transfer RNA (tRNA), enhancer RNA have been well documented to be decorated with many methylation marks such as 5-methylcytosine (5mC), N-6-methyl adenosine (m6A) and other marks (**1, 2, 3, 4**). Elucidation of the key proteins that deposit the methylation marks (writers: METTL3 and METTL14) for m6A, those that read the marks (readers: YTHDC1, 2, HNRNP etc.) and proteins that remove the marks (erasers: ALKBH5, FTO) are beginning to emerge (**5, 6, 7, 8**). Deposition of methylation marks have been shown to control mRNA stability, splicing, export and maturation of transcripts (**9**). RNA methylation has also been implicated in malignancies. For example, an eraser of m6A marks, ALKBH5, has been demonstrated in the maintenance of glioma stemness (**7**). Likewise, an eraser of m6A, FTO, has been implicated in acute myeloid leukaemia. Recently, it has been demonstrated that m6A is oncogenic in glioblastoma (**4**).

The epigenetic mark, 5-methylcytosine (5mC), is deposited both on DNA and RNA. While information is available for 5mC DNA methyltransferase (DNMTs), it has been shown that 5mC is deposited on the RNA by a group of RNA methyltransferases called NSUNs (**3**). These enzymes belong to the Nol1P/Nop2/SUN domain group of enzymes (**3**). They are responsible for depositing 5mC marks on mRNA, tRNA, vault RNA, ribosomes and enhancer RNA (**10**). The enzymes that comprise this group are NSUN1, NSUN2, NSUN3, NSUN4, NSUN5, NSUN6 and NSUN7 (**3, 10**). They have been demonstrated to methylate the residues of C72 of tRNA^**cys**^ and tRNA^**thr**^ (**3**). A recent genome wide study has expanded the repertoire of tRNA residues that are deposited with 5mC to include C48, C49 and C50 (**11**). NSUN6 cellular location has been demonstrated to be cytoplasmic. A UV-crossed linked study followed by sequencing has shown both protein coding, rRNA, miRNA, tRNA and mRNA are all deposited with the 5mC mark by NSUN6 (**3**). While details are beginning to emerge on the normal function of NSUN - mediated deposition of 5mC on normal cells, there is no information on how NSUN6 - mediated deposition of 5mC is involved in formation of tumors, maintenance of tumor stemness, tumor invasiveness and tumor response to therapy.

Glioblastoma is the deadliest of all primary brain tumors. Current treatment options available for glioblastoma are limited to temozolomide, radiation and electric field treatment. In a landmark paper, the status of MGMT promoter methylation has been shown to predict favorable response to temozolomide or not in about 45% of glioblastoma patients, while in 55% of the patients, the methylation status of MGMT promoter, does not predict response to temozolomide (**12**). To understand in an unbiased manner what genes control glioblastoma response to temozolomide (TMZ) (**13**), we performed a genome wide scale CRISPR knockout screen under the selection of temozolomide, to understand genes that control glioblastoma response to temozolomide. Combining the most enriched genes from the CRISPR screen with gene expression data from glioblastoma patients (n=631) in the cancer genome atlas (TCGA) revealed the RNA methyltransferase NSUN6 as one of the genes whose G-CIMP methylator status changes significantly (Welch t-test, p=0.0001) compared to the non G-CIMP group.

Here, we show that the RNA methyltransferase NSUN6 is a 5mC RNA specific methyl transferase on total and small RNA. Loss of NSUN6 confers resistance to TMZ in glioma cells both in gene edited and in TMZ resistant patient derived glioma xenografts. Mechanistically, our data show that NSUN6 through deposition of 5mC controls glioma cell response to TMZ by controlling transcriptional pausing via accumulation of NELFB and the RPS6KB2. The findings reveal that 5mC can control the rate of transcription and translation initiation elongation. Finally, we show that high expression of NSUN6 confers survival benefit to glioblastoma and in cancers such as bladder cancer, cervical cancer, oesophageal cancer, head and neck squamous cancer, testicular germ cell tumor, hepatocellular carcinoma, lung adenocarcinoma, lung squamous carcinoma and pancreatic ductal carcinoma. Our findings indicate that NSUN6 deposition of 5mC on cancer cells promotes transcriptional initiation and enhances translational initiation response to therapy, thus opening avenues to (1) use 5mC to control transcriptional pause and (2) modulate NSUN6 to promote tumor cells to respond effectively to therapy.

## Results

### Genome scale CRISPR and TCGA gene expression analysis reveal NSUN6 controls glioblastoma response to Temozolomide

To understand genes that controls glioblastoma response to the alkylating agent temozolomide (TMZ), we performed a genome scale CRISPR KO screen under TMZ selection (**Fig. 1A**). Our results revealed the hits that were preferentially enriched (p<0.05) as genes that controls glioblastoma susceptibility to TMZ (**Fig. 1B**). To understand the relevance of the enriched CRISPR hits in the context of the gene expression of the glioblastoma patients. We parsed all the enriched hits (p<0.05) from the CRISPR through the gene expression of glioblastoma TCGA data using the XENA software. We asked the question, do variables such as G-CIMP status affect the genes identified from the CRISPR. We found that the gene NSUN6, an RNA methyltransferase was the most enriched in the glioblastoma patients that changes significantly upon G-CIMP status (p=0.0001, Welch test, **Fig. 1C**). NSUN6 is an RNA methyltransferase (Haag S, et al RNA 2015), with little or no information of how it’s involved in cancer nor in chemotherapy. To understand, how NSUN6 might contribute in regulating susceptibility to TMZ in glioblastoma, we obtained six glioblastoma patient derived xenografts (**Fig. 2A)** and treated it with temozolomide (50-400µM) for 72hrs. We found that MES83 was the most susceptible to TMZ (**Fig. 2A**). Elsewhere, we have also evidence that U251, a clonal variant of SNB19 used in the CRISPR screen (**Fig. 1A**) is as susceptible to TMZ as MES83 which belongs to the same mesenchymal category of glioblastoma. To validate that NSUN6 contributes to TMZ susceptibility, we used the two guides that are enriched for NSUN6 under TMZ selection when compared with DMSO from the CRISPR screen (**Fig. 2B**). Using these guides, we performed a single gene editing of NSUN6 in MES83 and U251 (**Fig. 2C**) as previously described (Awah CU et al Oncogene 2020). Gene edit were confirmed by protein expression loss by western blot (**Fig. 2C, D**). We found that loss of NSUN6 conferred resistance to susceptible glioma cell lines (MES83, U251) (**Fig. 2E, F, G, and H**).

**Figure 1:**
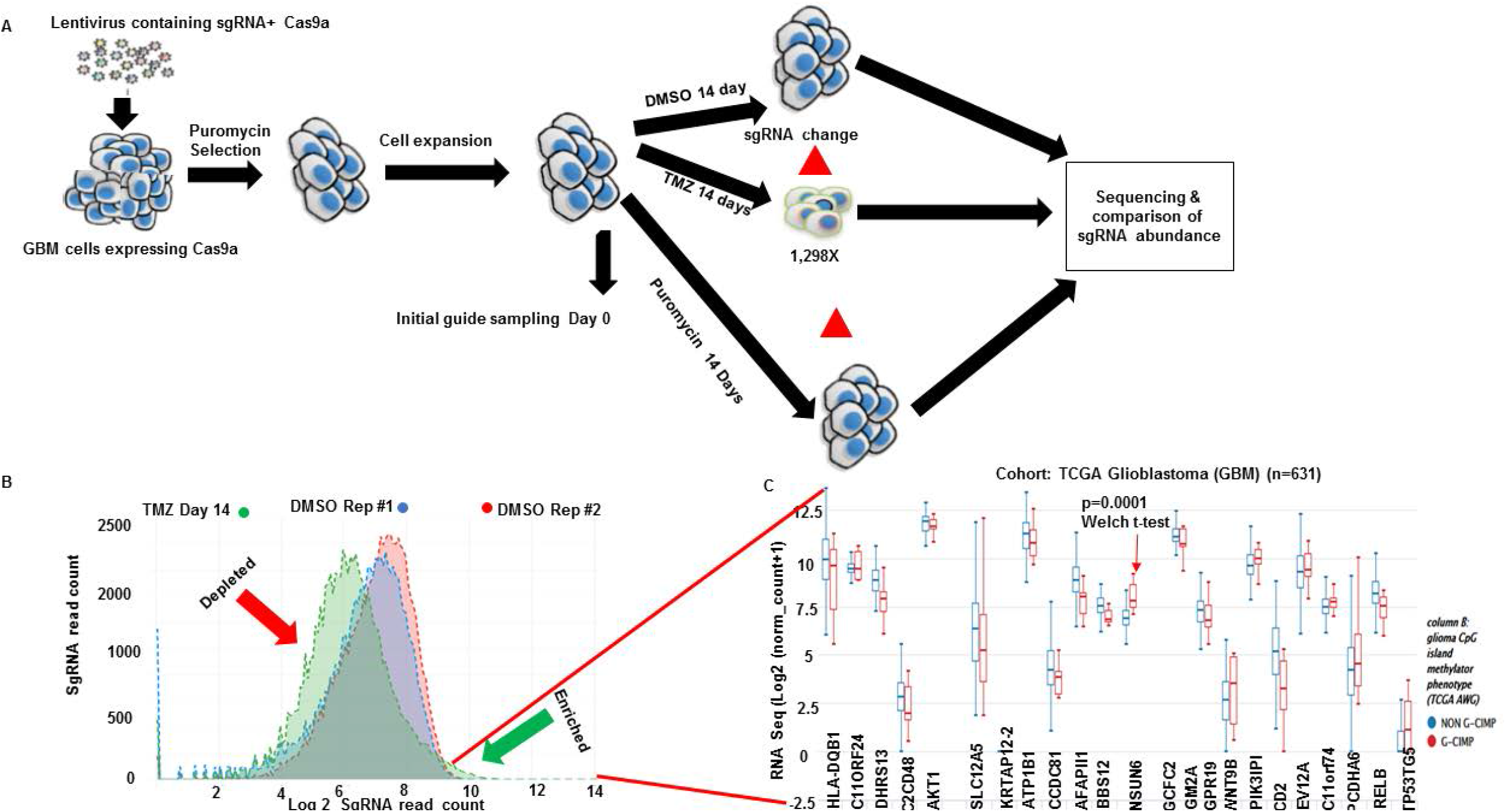
Genome wide CRISPR to reveals genes controlling response to Temozolomide (TMZ) **(A)** Schematic depiction of the CRISPR screen experiment performed in this study. Vector containing 76,441 sgRNA library expressing Cas9a were packaged into lentivirus, which was spinfected into SNB19 cells. The transfected cells were selected with puromycin for 96hrs. Cells were expanded and split into Temozolomide, DMSO and puromycin treatment for 14 days with 1×10^8^ cells per condition (1298X coverage; DMSO and puromycin were controls). Unique barcoded primers were used to amplify the library, puromycin/Day 0, DMSO day 14, Temozolomide day 14 and Puromycin day14 and selected guides. These samples were pooled and sequenced. SgRNA enrichment was analyzed using CRISPRAnalyzer (Winter J, 2017). **(B)** Distribution curve shows log2 fold read count for genes that were enriched in the temozolomide (green arrow) and those that were enriched in DMSO. **(C)** Bar charts show genes that were enriched in temozolomide (p=0.005) parsed through gene expression of TCGA glioblastoma patients with NSUN6 being the enriched in GCIMP status (p=0.0001, Welch t-test).

**Figure 2:**
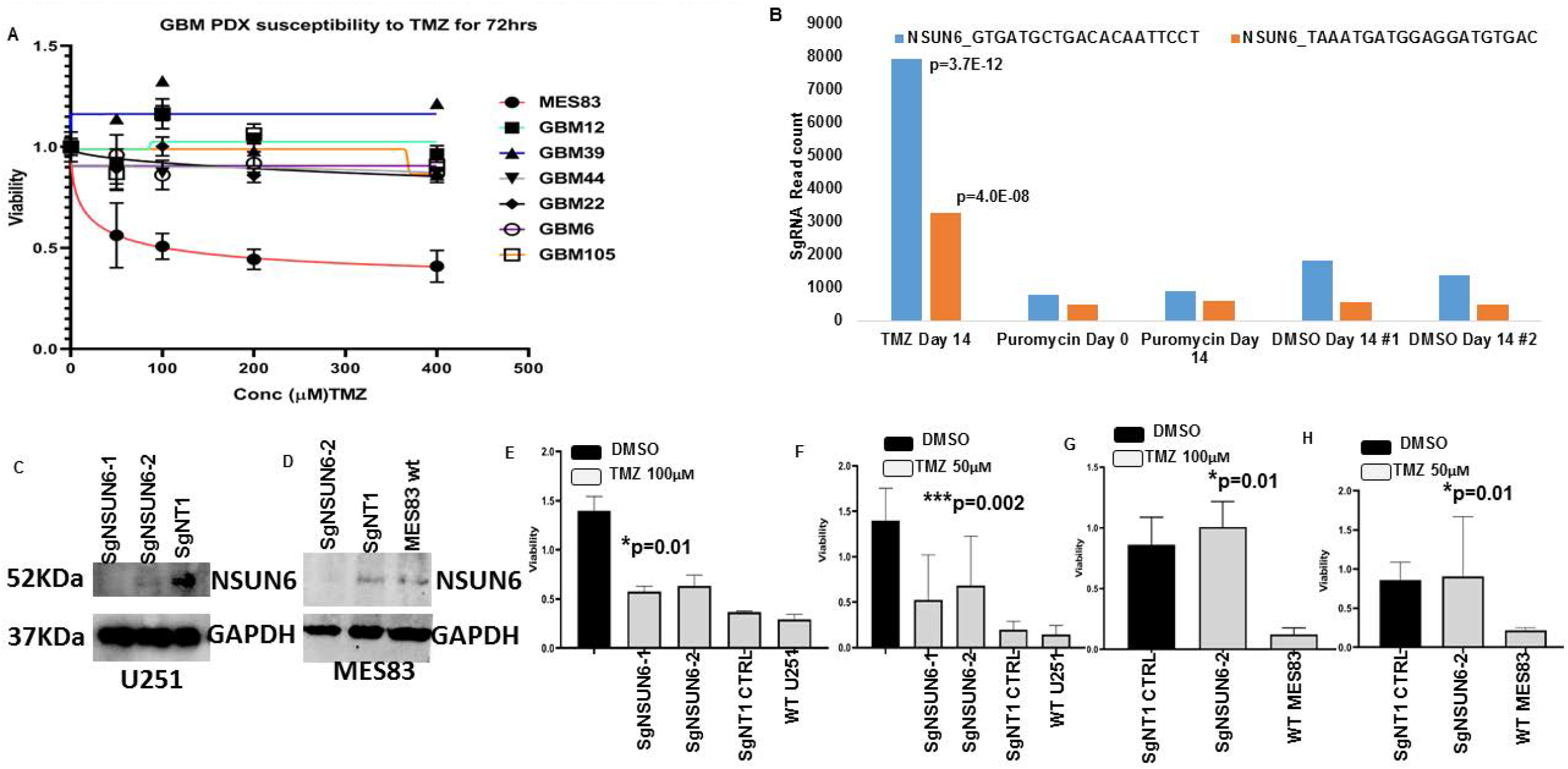
Genome wide CRISPR to reveals genes controlling response to Temozolomide (TMZ) **(A)** Temozolomide dose response curves of 7 human glioma cell lines treated with Temozolomide (50-400µM) and DMSO for 72hrs. (**B**) Bar chart shows the most enriched guides for NSUN6 (blue, orange compared in TMZ, Puromycin days 0, 14 and DMSO day 14 (control)(Rep#1,2). (**C&D**) Western blots show edited NSUN6 in U251 and MES83 compared to non-targeting controls & wild type. (**E, F, G & H**). Bar chart shows viability for SgNSUN6 edited cells and non-edited treated with TMZ (50µM, 100µM) for 72hrs.

### NSUN6 is a specific 5-mC RNA methyltransferase for large and small RNA in gliomas

NSUN6 has been demonstrated to be an RNA methyltransferase of tRNA in normal cells. We wanted to understand if the NSUN6 is expressed in glioma and what is the cellular localization of its signal in a malignant glioma. We performed immunofluorescence for NSUN6 and its methylation mark-5mC on patient derived glioma xenografts (MES83, GBM6, GBM44, and GBM39), we found that NSUN6 and 5mC signal is localized to the cytoplasm (**Fig. S1A, B**). To validate NSUN6 as a specific RNA methyltransferase in glioblastoma, we assayed for 5mC in sgNSUN6 edited U251 cells, we found loss of 5mC in the NSUN6 edited cells (**Fig. 3A**). Further investigation revealed loss of 5mC on both the total RNA and small RNA compared to wild type and non-targeting controls (**Fig. 3A, B, C, D, and F**). We treated NSUN6 edited RNA with RNASE and could show that 5mC is lost upon this treatment (**Fig. 3E**). Previous finding has specifically focused on the impact of NSUN enzymes on the methylation of tRNA residues (Haag S, et al RNA 2015; Huang T et al, NSMB 2019), our findings here reveals that NSUN6 impacts both large and small RNA to control glioma response to therapy.

**Figure 3:**
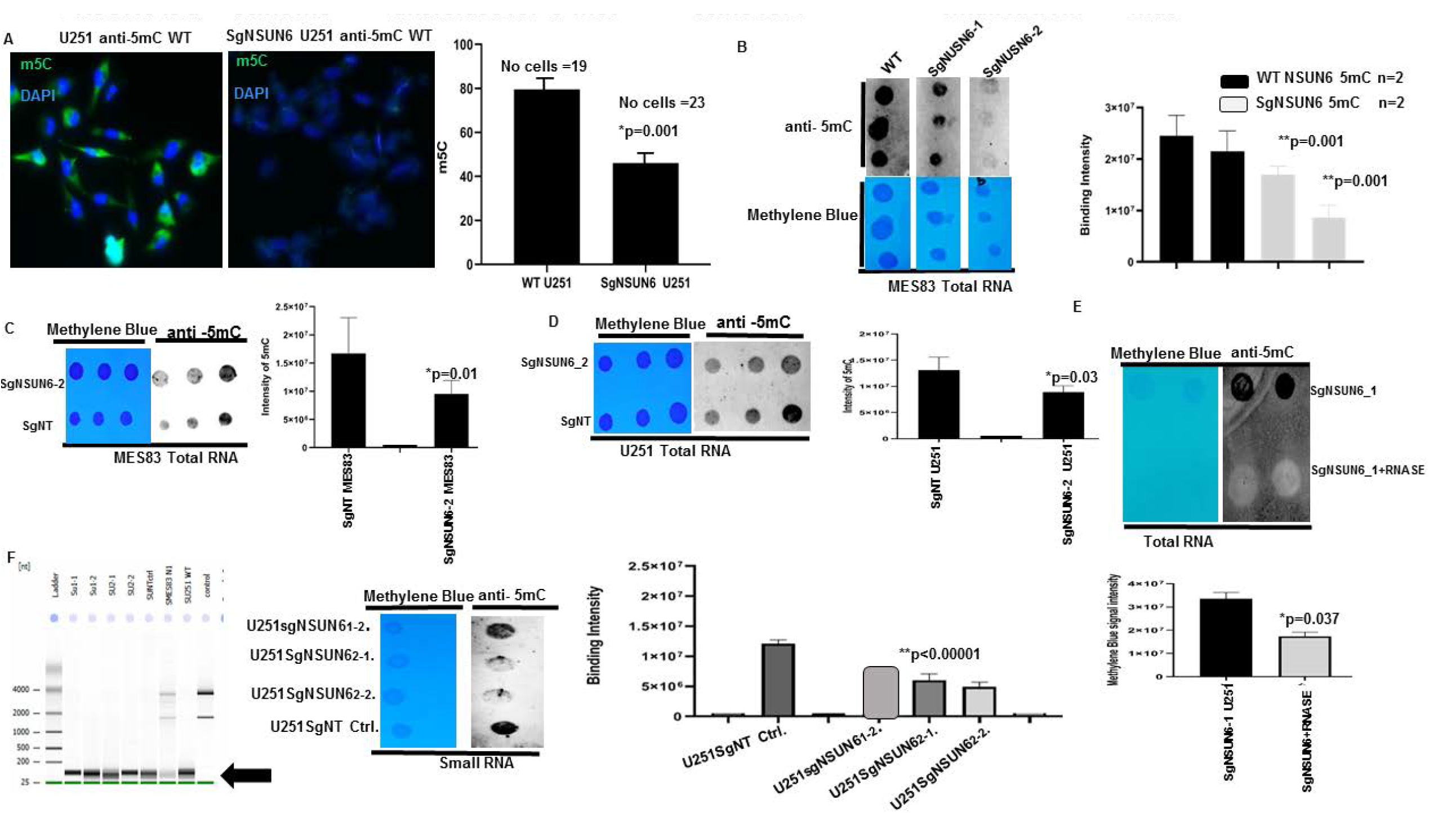
NSUN6 is a specific methyltransferase for 5-methyl cytosine for large RNA & small RNA in gliomas. **(A)** Picture shows 5mC staining for SgNSUN6 edited cells and wild type cells, intensities quantified in bars. **(B)** Dot blot shows MES83 total RNA spotting for SgNSUN6 (guide 1 & 2) and the wild type stained with methylene blue (below) and anti-5mC (upper), quantified in bar chart. (**C &D**) Dot blot shows spotting of total RNA of SgNSUN6 (guide 2) and SgNT for U251 and MES83 stained with methylene blue and anti-5mC and. **(E)** Blot shows dot blot of total RNA treated with and without RNAase and stained with methylene blue and anti-5mC and quantified below. (**F**) Gel picture shows small RNA extracted from edited cells and controls, dot-blotted and stained for methylene blue and anti 5mC and quantified.

### NSUN6 and 5-mC is lost in generated temozolomide resistant glioblastoma patient derived xenografts

Resistance to chemotherapy is a major concern in cancer therapeutics, we hypothesized that if NSUN6 and 5-mC is indeed a major contributor of glioma resistance to TMZ, then its expression should be lost on the TMZ resistant glioblastoma patient derived xenograft. To do this, we used GBM6 resistant to TMZ (GMB6R: GBM6 cells were implanted in mouse brain and orthopically treated with TMZ till they are very resistant). We extracted RNA from these cell lines comparing the untreated, DMSO day4, TMZ day4, GBM6 R, DMSO day 8 and TMZ day 8 and performed a dot-blot against 5mC on the RNA. We found that in the GBM6R and TMZ day 8 treated cells, there is loss of the 5mC marks (**Fig. 4A**). To investigate if the loss of 5mC is due to loss of NSUN6 expression on these cell lines. We performed dot bot, western blot on the protein extracts and found that the expression of NSUN6 is significantly lost on the cell lines that are resistant to TMZ: GBM6R TMZ and TMZ day 8 (**Fig. 4B, C**) and in a time dependent manner (**Fig. 4D**).

**Figure 4:**
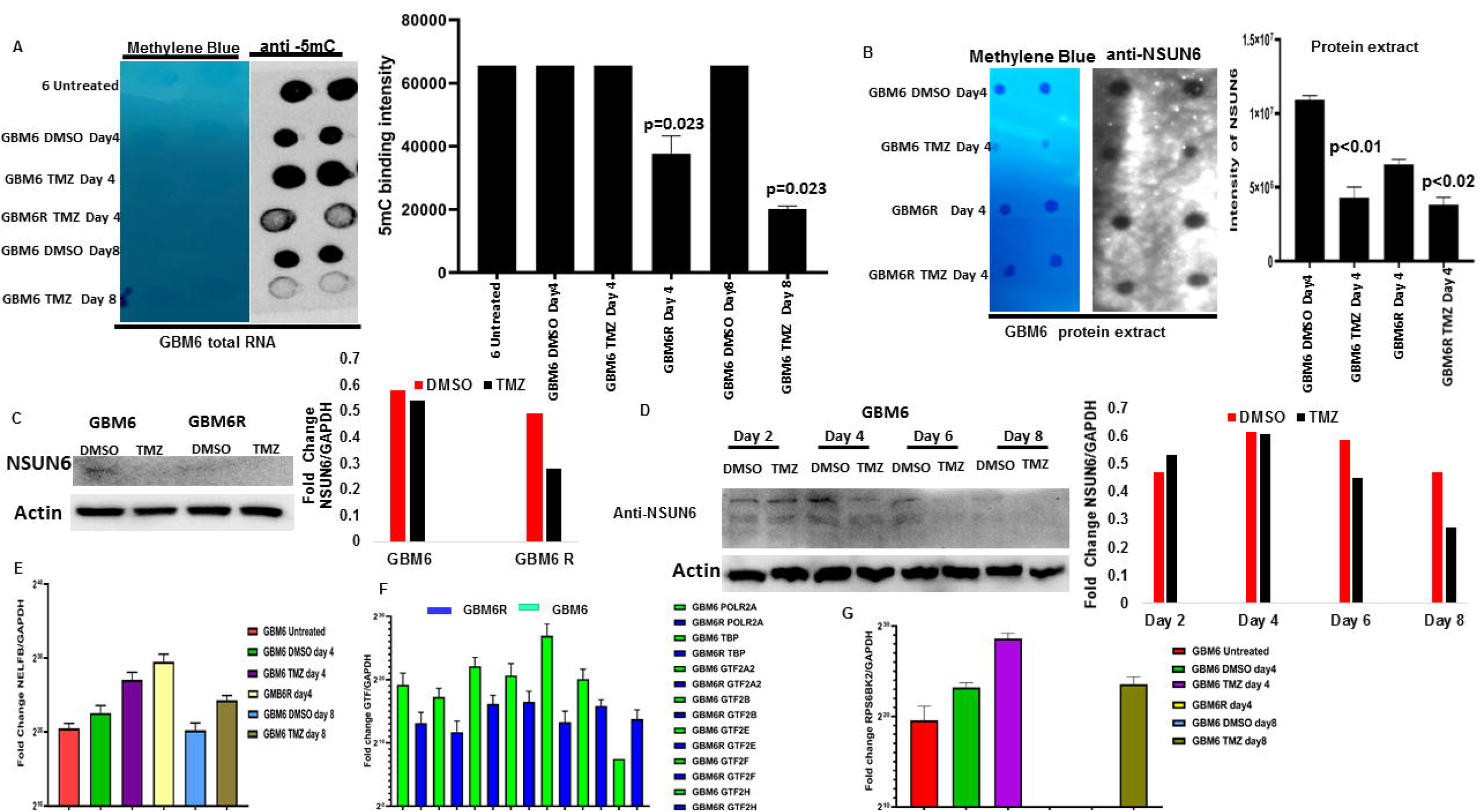
NSUN6 & 5-methyl cytosine expression decreases in generated TMZ resistant gliomas PDX xenografts. **(A)** Dot blot of total RNA shows GBM6 untreated, GBM6 treated with DMSO day4, GBM6 TMZ day 4, GBM6R TMZ day 4, GBM6 DMSO day 8, GBM6 TMZ day8 stained with methylene blue and anti-5mC, quantified on the right **(B)** Dot blot of protein extract of GBM6 treated with DMSO day4, GBM6 TMZ day 4, GBM6R TMZ day 4 stained with methylene blue and anti-NSUN6 and quantified on the right. **(C)** Western blot against NSUN6 and Actin B on GBM6 and GBM6R treated with TMZ and DMSO, quantified on the right. **(D)** Western blot against NSUN6 and Actin B treated DMSO or TMZ on day 2, 4, 6 and 8 and quantified on the right **(E)** Bar charts shows fold change enrichment of NELFB against GAPDH on GBM6 untreated, GBM6 treated with DMSO day4, GBM6 TMZ day 4, GBM6R TMZ day 4, GBM6 DMSO day 8, GBM6 TMZ day8.**(F)** Bar plots shows the fold change of POLR2A,TBP,GTF2A2,GTF2B, GTF2E, GTF2F, GTF2H normalized against GAPDH in GBM6 TMZ and GBM6R **(G)** Bar charts shows fold change enrichment of RPS6KB2 against GAPDH on GBM6 untreated, GBM6 treated with DMSO day4, GBM6 TMZ day 4, GBM6R TMZ day 4, GBM6 DMSO day 8, GBM6 TMZ day8.

### NSUN6 via 5-mC deposition regulates mRNA stability in time dependent manner and Temozolomide response through NELFB coordinated transcriptional pausing

Having established that the NSUN6 and 5mC is implicated in the temozolomide resistant phenotype found in the GBM6R glioma cell lines. We wanted to understand in mechanistic terms how NSUN6 regulates this phenotype. We treated the NSUN6 and WT U251 cell lines with Actinomycin D in time dependent manner as described (**15**) and assayed for 5mC marks and total RNA stability by one phase decay kinetics. We found that 5mC decreases in the WT cells with maximum decrease in 5mC signals at the 24hr (**Fig. 5A**), conversely the 5mC signals is lost at 2hrs and 8hrs on the NSUN6 edited cells but accumulates at the 24hrs (**Fig. 5B**). This implies that (i) 5mC is involved in transcription and mRNA translation, that once transcription is blocked, nascent mRNA losses 5mC marks (ii) that NSUN6 is a major RNA methyltransferase but not the exclusive RNA methyltransferase in glioma. With 5mC loss upon blocking of transcription and decrease in mRNA, we reasoned that NSUN6 via 5mC might couple transcription to translation. To find the genes through which this process might be mediated, we analyzed the genome scale NSUNs 5mC methylated genes already identified (**11**). We examined for NSUN 5mC target genes that overlapped between in the human cortex (brain), and HEK293T (**Fig. 5C**). Since the human experimentally validated genes in this data is that of those overlapped between human brain cortex and HEK293T cells. In this subset, we found about 109 genes that were significant (p<0.0000001) in this group (**Fig. 5C**). To further investigate this subset, we parsed them through our genome scale CRISPR TMZ screen and searched for the genes in our both enriched and depleted datasets (**Fig. 5D**). Of these NSUN 5mC methylated genes, 8 genes were (enriched genes in TMZ) and 10 genes (depleted in TMZ) (**Fig. 5D**). Of these genes, since we are interested in exploring the link on how NSUN6 couples transcription and translation to regulate TMZ response, the NELFB and the RPS6KB2 were attractive targets. NELFB has been extensively studied (**20**) as a being involved in stalling RNA POLII on the paused and arrested transcriptional complexes. Having identified NELFB and RPS6KB2 as one of the genes involved the NSUN6 mediated deposition of 5mC as well as implicated on our TMZ CRISPR screen, then we reasoned that as they might be key players in transcriptional pausing as well as translation initiation, that if they are impacted by loss of NSUN6 and associated 5mC mark, then we should see changes in their mRNA stability upon inhibition with Actinomycin D which blocks both RNAPOLI & II. Indeed, we found that loss NSUN6 (**Fig. 5E & 5G**), lead to the delay in the decay of the NELFB and RPS6KB2, as well as in TBP & POLR2A (**Fig. S2A & B**) which could be interpreted as accumulation of the NELFB, RPS6KB2 and preinitiation GTFs on the NSUN6 edited cells. To corroborate this finding in the GBM6R model, described in **Fig. 4**. We analyzed for the expression levels of NELFB and RPS6KB2 in the GBM6R, we found in (**Fig. 4E**) that NELFB accumulates on the GBM6 TMZ day 4 compared to DMSO day 4 and on the TMZ day 8 compared to the DMSO Day8. The NELFB accumulates the highest on the GBM6R (**Fig. 4E**). For the RPS6KB2, we found that it accumulates on the GBM6 TMZ day 4 and GMB6 TMZ day 8, we found no RPS6KB2 on the GBM6R and the GBM6 DMSO day 8 (**Fig.4G**). With NELFB and RPS6KB2 found to be accumulating in GBM6 resistant to TMZ and NSUN6 edited cells (**Fig. 4E, G & Fig. 5E, F, G, I**). We reasoned that if NSUN6 via 5mC controlled NELFB and RPS6KB2 transcript, then an immunoprecipitation of 5mC from NSUN6 edited cells and the GBM6 R cells and subsequent analysis of the transcript of NELFB and RPS6KB2 should show an accumulation of NELFB and RPS6KB2. Indeed, we found that loss NSUN6 and 5mC lead to accumulation of NELFB and RPS6KB2. Using 5’deoxy-2’aza-cytidine, we treated wildtype gliomas and assayed for the loss of 5mC and the expression of GTF, we found that NELFB is increased in the azacytidine treated cells compared to the other GTF (**Fig. S2C, D**). The findings that NELFB accumulates in the absence of NSUN6 and 5mC points to the fact that NSUN6 via 5mC is regulating transcriptional pausing.

**Figure 5:**
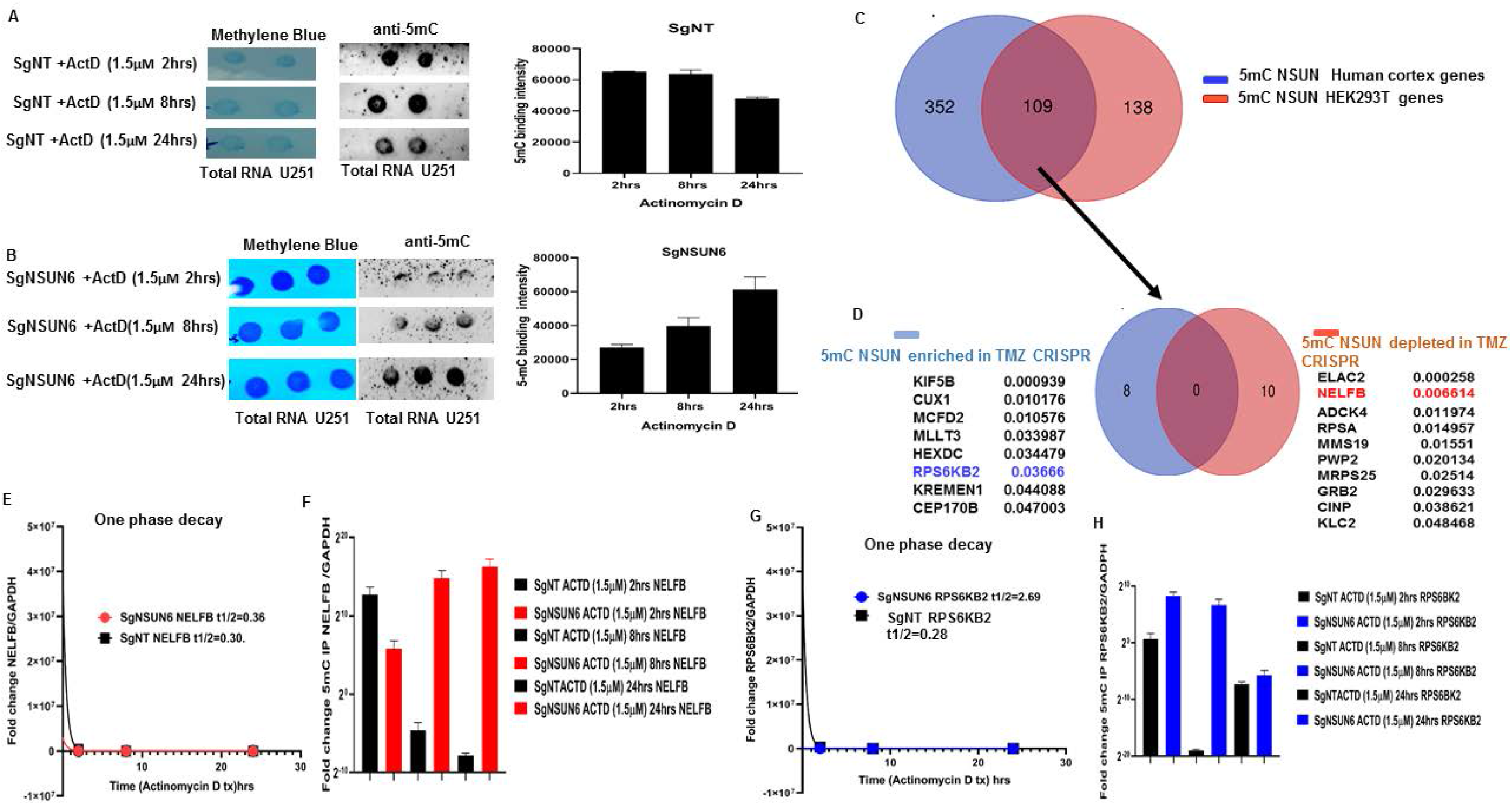
NSUN6 via 5mC deposition regulates mRNA stability in time and transcriptional dependent manner via NELFB/RPS6KB2. **(A)** Dot blot of total RNA of SgNT U251 treated actinomycin D at 2hrs, 8hrs and 24hrs and stained with methylene blue and anti-5mC and quantified on the right. **(B)** Dot blot of total RNA of SgNSUN6 U251 edited cells treated with actinomycin D at 2hrs, 8hrs and 24hrs and stained with methylene blue and anti-5mC and quantified on the right. **(C)** Venn diagram represent gene that are methylated by NSUN in human brain cortex and HEK293T cells with 109 genes that overlapped (Huang T et al, NSMB 2019). **(D)** Venn diagram represents 109 NSUN methylated genes that were overlapped between human brain cortex and HEK293T that are enriched in CRISPR TMZ (blue) and depleted in TMZ (brown). **(E)** Graph shows one phase decay half life curve comparing NELFB mRNA stability in SgNT (black) and SgNSUN6 (red) U251 total RNA. **(F)** Bar charts represents fold change of 5mC IP enrichment of NELFB normalized against GAPDH comparing SgNT (black) versus SgNSUN6 (red) total RNA. **(G)** Graph shows one phase decay half-life curve comparing RPS6KB2 mRNA stability in SgNT (black) and SgNSUN6 (blue) U251 total RNA. **(H)** Bar charts represents fold change of 5mC IP enrichment of RPS6KB2 normalized against GAPDH comparing SgNT (black) versus SgNSUN6 (blue) total RNA.

### TATA binding protein (TBP) complexes formation and general transcription factors (GTF) accumulates on the preinitiation sites in the absence of NSUN6 and 5mC

Since, NELFB is involved in transcriptional pausing, we wanted to explore at what point in transcription mechanism does NELFB accumulates in the absence of NSUN6 and 5mC and in GBM6R TMZ phenotype. We obtained TBP probe (**19**) and performed a DNASE footprinting assay as previously described (**14, 16, and 17**) with the NSUN6 edited and non-target control edited cells with and without Sarkosyl (an inhibitor of NELFB). We found that (**Fig. 6A**) three complexes were formed as also seen elsewhere previously (**21**). Complexes I, II & III are seen on the lane 2 for NT control but complex II is not seen on lane 1 for NSUN6 edited cells indicating that TATA binding proteins complexes is bound on that site and thus could not be cleaved by DNASE 1 and this supports our finding that transcription pre-initiation complexes accumulates in the absence of NSUN6 and 5mC as shown in **Fig. 4E, G** and **Fig. 5E, F**. To investigate what is the expression profiles of the general transcription factors complexes (GTFs) which are TBP, POLR2A, GTF2A2, GTF2B, GTF2E (**21, 23**). We performed an 5mC IP on the sgNSUN6 cells and NT controls treated with Actinomycin D in a time dependent manner as well on the GBM6 and GBM6 R TMZ treated cells and probed for the expression of the TBP, POLR2A, GTF2A2, GTF2B and GTF2E and found that TBP, POLR2A, GTF2A2 and GTF2E (**Fig. 6B, C, D & F**) is enriched in the NSUN6 edited cells. Subsequently, we analyzed for the enrichment of GTF in GBM6 versus the GBM6R, we found that POLR2A, TBP, GTF2A2, GTF2B, GTF2E, GTF2F were enriched in GBM6 (**Fig. 4F**) compared to GBM6R.

**Figure 6:**
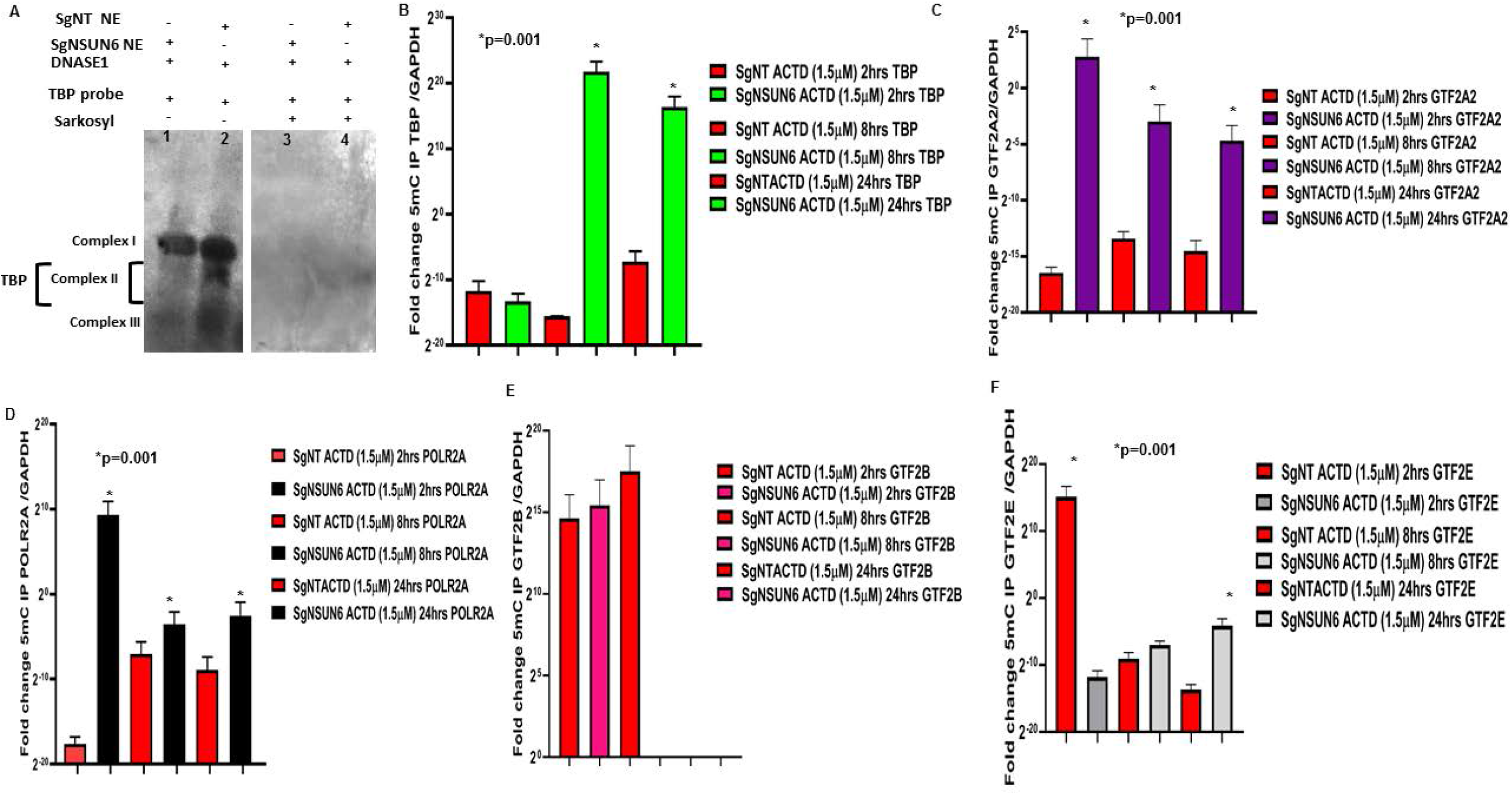
NSUN6 via 5mC regulates TATA binding protein complexes formation and the preinitiation complex formation. **(A)** Blot shows TATA binding protein protein (TBP) DNase foot printing in SgNT (lane 1) and SgNSUN6 (lane 2) and lane (3) SgNT treated with sarkosyl and lane (4) SgNSUN6 treated with sarkosyl. **(B)** Bar charts represents fold change of 5mC IP enrichment of TATA binding protein (TBP) normalized against GAPDH comparing SgNT (red) versus SgNSUN6 (green) total RNA. **(C)** Bar charts represents fold change of 5mC IP enrichment of GTF2A2 normalized against GAPDH comparing SgNT (red) versus SgNSUN6 (purple) total RNA **(D)** Bar charts represents fold change of 5mC IP enrichment of POLR2A normalized against GAPDH comparing SgNT (red) versus SgNSUN6 (black) total RNA. **(E)** Bar charts represents fold change of 5mC IP enrichment of GTF2B normalized against GAPDH comparing SgNT (red) versus SgNSUN6 (magenta) total RNA. **(F)** Bar charts represents fold change of 5mC IP enrichment of GTF2E normalized against GAPDH comparing SgNT (red) versus SgNSUN6 (grey) total RNA.

### High NSUN6 expression confers survival benefits to glioblastoma and other cancers

To explore if the high expression of NSUN6 confers survival benefit to glioblastoma patients or not. We first ruled out the impact of MGMT promoter status on the influence of NSUN6 survival benefit, if there is any (**12**), because MGMT promoter status has already been established as a predictor of glioblastoma response to temozolomide. We ran a multivariate correlation analysis comparing if the expression of MGMT impacted NSUN6 and vice versa. We found that the expression of MGMT does not have any impact on the NSUN6 (**Fig. S3A**). Having established that there is no impact of MGMT on NSUN6, then we explored if high expression of NSUN6 conferred favorable survival benefit or not to glioblastoma patients. We found across different TCGA GBM datasets analyzed with GlioVis (**24**) that high expression of NSUN6 conferred survival benefit to glioblastoma patients (**Fig. 7A**). Next, we explored if the MGMT promoter status of the glioblastoma patients impacted their survival outcome, we found that MGMT promoter status had no impact on the NSUN6 based survival outcome of the patients (**Fig. 7B**). That is with MGMT promoter status or not, the NSUN6 expression conferred a favorable survival status to glioblastoma patients who have high expression of NSUN6 thereof. To extend this finding, we used Kaplan-Meier plotter (**25**) and analyzed different cancers, if the high expression of NSUN6 conferred survival benefit or not. Indeed, we found that high expression of NSUN6 in cancer such as bladder cancer, cervical squamous carcinoma, esophageal squamous carcinoma, head & neck squamous carcinoma, testicular germ cell tumor, hepatocellular carcinoma, lung adenocarcinoma, lung squamous carcinoma, pancreatic ductal carcinoma (**Fig. 8A, B**) conferred favorable survival outcome. This indicates that high NSUN6 expression might be a prognostic marker for these tumors.

**Figure 7:**
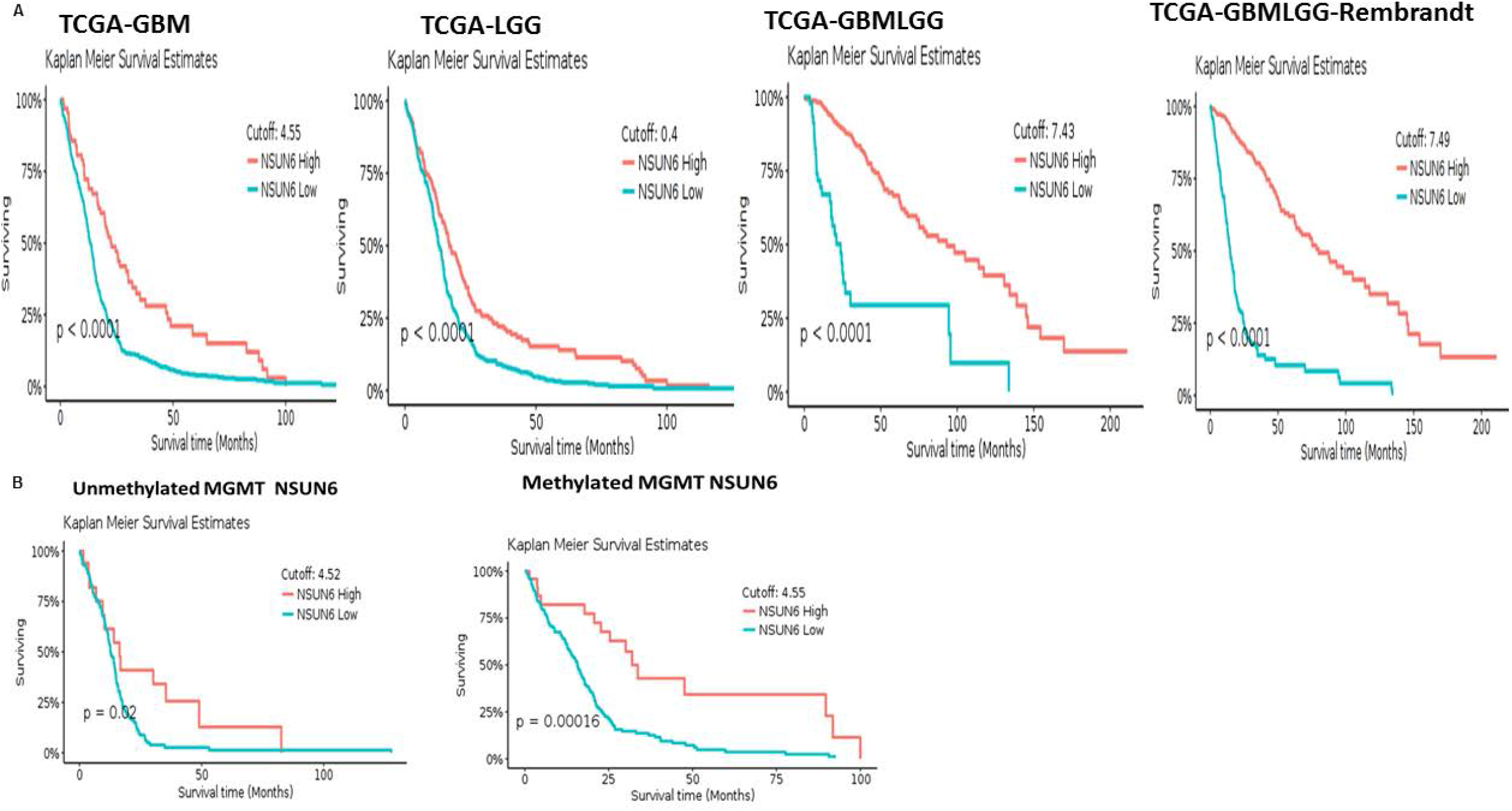
NSUN6 expression confers survival benefit to GBM and LGG patients treated with/out MGMT promoter methylation status. **(A)** Kaplan Meier survival curve shows high expression of NSUN6 versus low expression across TCGA GBM data set **(B)** Kaplan Meier survival curve comparing high versus low expression of NSUN6 in methylated and unmethylated TCGA GBM patients.

**Figure 8:**
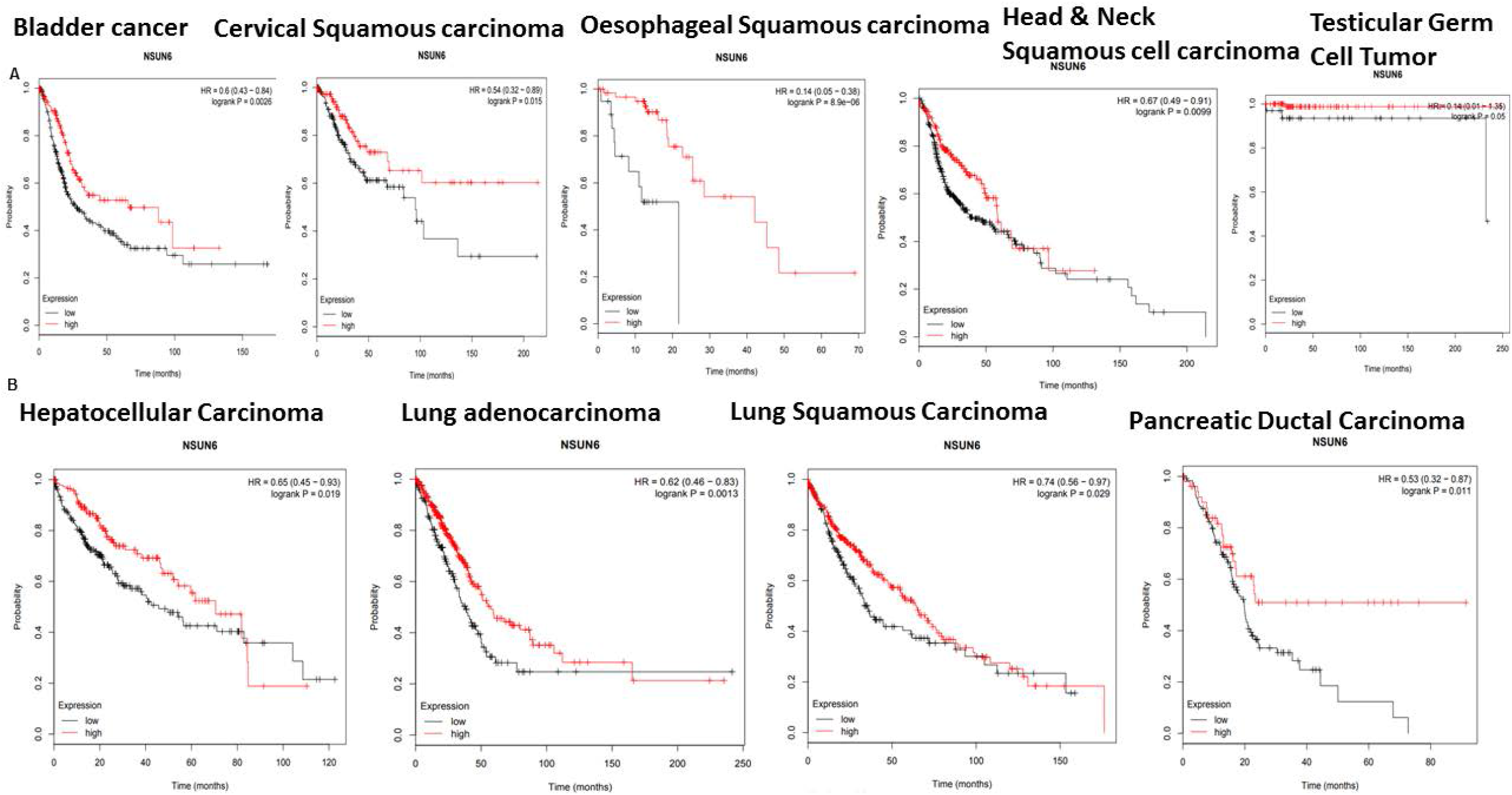
High NSUN6 expression confers survival benefit to other cancers. **(A&B)** Kaplan Meier survival curve compares high versus low expression of NSUN6 across cancers (bladder cancer, cervical squamous cancer, oesophageal cancer, head & neck cancer, testicular germ cell tumor, hepatocellular cancer, lung adenocarcinoma, lung squamous carcinoma and pancreatic ductal carcinoma.

## Discussion

Epitranscriptomics is an emerging field revealing a new layer of gene regulation. Various RNA molecules such as mRNA, tRNA, enhancer RNA, vault RNA and microRNA have been shown to be decorated with many marks such as 5mC, m6A, m6m etc. Whilst m6A has been explored in cancers (**6, 7, 8**). It is not known the impact of NSUN6 in particular in relations to oncogenesis and tumor response to therapy. Various NSUN such as NSUN2 and NSUN3 has been linked to cancers like leukemia, precisely NSUN3 has been linked to 5-azacytidine sensitivity to leukaemia through P-TEFB recruitment of RNAPOLII (**26**). NSUN2 interaction with METTL has been shown to sensitize Hela cells to 5-fluorouracil (**27**). In-addition, NSUN2 has been shown to interact with the proto-oncogene MYC to drive cellular proliferations in tumors (**28**). Moreso, high expressing NSUN2 patients has been shown to have superior overall and disease progression free survival in ovarian cancer (**29**). Here-in, we have demonstrated that NSUN6 deposits 5mC on glioblastoma to control response to therapy via regulation of the transcriptional pausing through NELFB and that high expression of NSUN6 across cancers is prognostic of a favorable survival outcome.

RNAPOLII are known to be poised, paused or arrested in relation to gene expression states and environmental cues during transcription (**20**). RNA POLII pausing has been posited to be paused or arrested in cellular stress yet the mechanistics of this is yet lacking (**30**). Bugai A et al demonstrates that RBM7 mediated recruitment of RNAPOLII and P-TEFB drives transcriptional activation in response to genotoxic DNA damage induced stress. In-contrast, our works shows the opposite, in that our findings points to a model where in NSUN6 is required to deposit 5mC on the NELFB, thus releasing transcription pausing at the pre-initiation complex, which will allow the recruitment of the general transcription factors and RNAPOLII and TBP for transcription to proceed. This mechanism is impaired in cells that are NSUN6 edited. Moreso, we present evidence that supports this finding in a GBM6R TMZ resistant model, where we show that there is accumulation of NELFB, with reduction of other general transcription factor when there is loss of NSUN6 and 5mC. Again, this finding is in support of what is observed for NSUN3, in which NSUN3 mediated deposition of 5mC sensitizes the recruitment of the positive transcription elongation factors P-TEFB to drive RNAPOLII with cells responding to treatment.

The results in SgNSUN6 edited cells and the GBM6R points to a general mechanism, where in NSUN6 deposit 5mC on the NELFB and releases the break on transcriptional pausing during cellular stress that occurs during treatment with the alkylating agent temozolomide. This finding together with that of Cheng X et al, 2018 points to a novel role of NSUNs, in particular NSUN6 and NSUN3 involvement in regulation of RNAPOLII based transcription as well as offer a novel mechanistic insight into how transcription is regulated in states of cellular stress. It is plausible as there has been many efforts into understanding how transcription can be controlled in time and space, that our findings points to a possible universal mechanism in which the transcription of a cell can be regulated both by the NSUN methyltransferases and the methylation marks they deposit, which is 5mC.

## Materials and Methods

### Genome wide scale CRISPR-Cas9a knockout screen

As previously described (**14**), we used the Brunello Library that contains 70,000 sgRNA which covers the 20,000 genes in the human genome at the coverage rate of 3-4sgRNA/gene plus 10,000 sgRNA which are non-targeting controls. To perform the CRISPR screening, glioma SNB19 cells were expanded to 500 million and then spininfected with 70,000sgRNA. After spinfection, the cells are selected with 0.6µg/ml of puromycin for 4 days. This selection is aimed at the cells that have been rightly integrated with the sgRNA that incorporates the puromycin cassette into their genome. We achieved an MOI of 21% in two independent screens. At the end of day 4, about 150 million cells survived the selection. We used 50 million selected cells for the extraction of genomic DNA. The base sgRNA representation is obtained by amplification of the sgRNA with unique barcoded primers. The remaining 100 million cells were expanded for 2 days, until cells grew to 300 million. 100 million cells each were treated with temozolomide at concentration of 700µM TMZ or DMSO or puromycin for 14 days. After 14 days, the cells were harvested, the gDNA extracted, and the sgRNA amplified with another unique barcoded primer.

### DNA extraction and PCR amplification of pooled sgRNA

As already described (Awah CU et al 2020 Oncogene). The genomic DNA (gDNA) were extracted with the Zymo Research Quick-DNA midiprep plus kit (Cat No: D4075). gDNA was further cleaned by precipitation with 100% ethanol with 1/10 volume 3M sodium acetate, PH 5.2 and 1:40 glycogen co-precipitant (Invitrogen Cat No: AM9515). The gDNA concentration were measured by Nano drop 2000 (Thermo Scientific). The PCR were set up as described. The sgRNA library, puromycin, DMSO and etoposide selected guide RNA were all barcoded with unique primers as previously described.

### Next Generation Sequencing

The sgRNAs were pooled together and sequenced in a Next generation sequencer (Next Seq) at 300 million reads for the four sgRNA pool aiming at 1,000reads/sgRNA. The samples were sequenced according to the Illumina user manual with 80 cycles of read 1 (forward) and 8 cycles of index 1. 20% PhiX were added on the Next Seq to improve library diversity and aiming for a coverage of >1000reads per SgRNA in the library.

### CRISPR screen data analysis

All data analysis was performed with the bioinformatics tool CRISPR Analyzer (**18**). Briefly, the sequence reads obtained from Next Seq were aligned with human genome in quality assessment to determine the percentage that maps to the total human genome. To set up the analysis, the sgRNA reads (library, puromycin, DMSO and etoposide) replicates were loaded unto the software. The sgRNA that does not meet a read count of 20 is removed. Hit calling from the CRISPR screen was done based on sgRSEA enriched, p<0.05 was used for significance based on Wilcoxon test.

### Data availability

All data underlying the results are available as part of the article and no additional source data are required.

**DOI**: 10.5281/zenodo.5167143

Data 1: sgRNA library

Data 3: DMSO treated sgRNA library replicate 1

Data 4: DMSO treated sgRNA library replicate 2

Data 5: Puromycin treated sgRNA Library replicate 1

Data 6: Puromycin treated sgRNA library replicate 2

Data 7: Temozolomide treated sgRNA library

### Immunofluorescence

GBM patient-derived xenograft (PDX) lines, MES83, U251, GBM6, GBM6R, GBM12 and GBM44, GBM12, GBM39, GBM22, GBM105 cells. Cells were fixed in 4% PFA for 10mins at 4°C. After which they were blocked with 10% BSA in PBS for 2hrs at room temperature. Cell were washed and then incubated with anti-NSUN6, and anti-5mC (1:1000) (abcam) overnight at room temperature. Tissue were blocked in 3% BSA in PBS and incubated for 2hrs. Using anti-rabbit Alexa 488 and DAPI mounting media, we stained the proteins and obtained images on confocal microscope.

### Single gene editing

To edit NSUN6, we used single guide RNAs that were enriched for gene NSUN6 as well as the non-targeting controls as described (Awah CU et al 2020 Oncogene). Briefly, these guides were synthesized by Synthego and following the protocol, we prepared the ribonucleoprotein complexes by mixing the guides (180pmol) with recombinant Cas9 protein (Synthego) 20pmol in 1:2 ratio. The complexes were allowed to form at room temp for 15mins, and then 125µL of Opti-MEM I reduced serum medium and 5µL of lipofectamine Cas9 plus reagent were then added. Both, the cells and the formed ribonucleoprotein complexes, were seeded at the same time with 150,000 SNB19 cells in a T25 flask. The cells were incubated for 4days. After 4 days, the cells were harvested, and downstream analysis were performed to prove the editing of the genes.

### Western blotting

To confirm the loss of protein expression of the gene of interest following editing, we extracted the proteins using M-PER (Thermoscientific: 78501) and cocktail of phosphatase and protease inhibitors. The cells were lysed using water bath ultrasonicator for 4mins. Cell lysate were cleared by centrifugation. We measured the concentration of protein in lysates. Denatured lysates were loaded into 4-20% Tris-glycine gels (Novex) and separated at 180V for 2hrs. The gels were transferred unto a PVDF membrane by semi-dry blotting for 1hr. We blocked the membrane in 5% non-fat milk TBST buffer for 30mins and incubated with primary antibodies NSUN6 (1:500), GAPDH or ACTB (1:1500) in 5% BSA respectively over night shaking at 4°C. Primary antibodies were removed and we added the secondary polyclonal HRP (1:20,000) in TBST and incubated shaking for 2hrs at room temp. The membrane were washed 6X in TBST and then developed with ECL (Cat No: 1705061) and band imaged on a Bio-Rad Chemi-doc imaging system.

### Dot blotting

Briefly, RNA were extracted with miRVana RNA extraction kit (Thermofisher). The RNA were nanodropped and then spotted on the hybond N+ membrane (GE Healthcare) as previously described (**17**). Equal amount of the RNA were spotted and then cross linked by UV radiation for 10mins. After which membrane were stained with methylene blue for 10mins. Methylene blue were rinsed off under running cold water, and the RNA image were taken and the blot were blocked in 10% milk in TBST for 1hr and after which 1:1000 anti-5mC (Abcam) in BSA were incubated with the blot shaking overnight. Primary antibodies were removed and we added the secondary polyclonal HRP (1:20,000) in TBST and incubated shaking for 2hrs at room temp. The membrane were washed 6X in TBST and then developed with ECL (Cat No: 1705061) and band imaged on a Bio-Rad Chemi-doc imaging system.

### Viability assays

The edited cells (NSUN6, non-targeting control and wild type unedited U251 and MES83) were seeded at 4,000 cells/well in a 96 well plate and treated them with 50µM & 100µM TMZ or DMSO for 72hrs. For GBM PDX lines, we seeded them as well at 4,000cells/well in a 96 well and then added etoposide at a range of 50-500µMfor 72hrs. Titre glo was added following incubation with drugs. (Cat No: G7572) and the viability of cells was analyzed 5 min later by measuring the luminescence. We normalized the intensity against DMSO treated cells of each cell line or PDX or the edited cell and then determined the survival.

### qRT-PCR (Quantitative Reverse Transcript Polymerase chain reaction)

The SgNSUN6 edited U251 cells as well as the non-targeting controls cells were treated with actinomycin D (1.5μM) for 2, 8, and 24hrs. The total RNA was extracted using MiRVana RNA extraction Kit (Thermofisher). The quality of the RNA was determined by RNA pico bioanalyzer measuring the 18S and 28S ribosome. Using the superscript III first strand synthesis system (Cat no: 18080-051), we generated cDNA and performed qPCR with NELFB, RPS6KB2, TBP, POLR2A, GTF2A2, GTF2B, GTF2F, GTF2E, GTF2G and GTF2H and GAPDH primers in triplicates and fold change of expression of genes of interest were normalized against GAPDH.

### DNAse footprinting assay

Briefly as previously described (**14**). We obtained nuclear extract of sgNSUN6 and non-targeting controls edited cells as described (**16**,**17**) and incubated it with TBP probe labelled with biotin on 5’ and 3’ end as described (**19**) and followed protocol described (**14**). To inhibit, the TBP complexes, we treated the nuclear extracts with Sarkosyl, an inhibitor of NELFB. We then resolved the resolved the complexes formed in 4-20% Tris Glycine (Novex) and blotted it unto a hybond N+ membrane. The membrane was air dried and then blocked with nucleic acid blocking reagent (Roche) in PBS for 1hr and then incubated with 1:1000 Streptavidin AP (Roche) for 2hrs and then washed with PBS and then imaged with CDP star (Roche) in a biorad chemidoc imaging system.

### mRNA stability assay

To determine if the loss NSUN6 impacted transcript stability, we treated cells (both edited and non-targeting controls) with Actinomycin D at 2hrs, 8hrs and 24hrs as described (**15**). We extracted total RNA with MiRVana RNA extraction kit and quantified and performed dot bot for 5mC as described above. Next, we performed RT-PCR using specific primers for NELFB, RPS6KB2, and the GTFs, the performed qPCR were normalized against GAPDH in triplicates.

To calculate the stability of the mRNA, we used the methods as previously described (Ratnadiwakara M & Anko ML, Bioprotocol 2018), the one phase decay model on Graphpad prism and calculated the half-life decay of the transcript comparing SgNSUN6 and SgNT edited cells.

### 5-Aza-2-deoxycytidine assay

As a complementary prove of the impact of 5mC on the general transcription factors, we treated the cells with 5-aza-2-deoxycytidine which generally demethylates the cells. We dissolved the AZA in acetic acid, first in 1:1 acetic and water and then 1:1 into the container with Azacytidine making a 5mg solution.

### Gliobastoma patient derived xenograft culture

The patient’s derived xenografts GBM12, GBM6, GBM83, and GBM44, GBM22, GBM39, GBM105, GBM157 were used in this study. Briefly, all the GBM PDX cells were all authenticated, they were cultured in 1% FBS in DMEM media. SNB19 were grown in 10% FBS in MEM media containing, essential amino acids, sodium pyruvate and 1% glutamine. U251 were grown in 10% FBS DMEM media. The cells were all grown to 80% confluency and then used for downstream analysis.

### Statistical analysis

CRISPR analysis were all performed with CRISR Analyzer (Winter J et al, 2017) which contains 8 statistical analysis for hit calling. All our experiments were performed in at least two independent experiments with multiple replicates. All bar charts in the manuscript were built with Graph Pad prism software 8 (San Diego, CA, USA). The statistical analysis performed for each figure are listed in the figure of the accompanying figures.

## Contributions

CUA conceived this project. CUA performed the CRISPR screen, single gene editing and all experiments, CUA prepared the SgRNA library for sequencing and analyzed the CRISPR screen with JW. CMM contributed to experiment. AS provided material. CUA performed all the validation experiments. CUA and OOO wrote the manuscript.

## Acknowledgements

The authors acknowledge the support of the members of labs at Northwestern Feinberg School of Medicine. The authors acknowledge the materials and reagent given for this work by the Adam Sonabend laboratory at the Northwestern University, Chicago. The authors thank Prof Kevin Struhl of the Harvard Medical School for the criticism of the data.

## Declaration

The authors declare no conflict of interest.

## Supplementary Figure Legends

**Supplementary Figure 1: Expression of NSUN6 correlates with its RNA methylation mark 5mC in the cytoplasm of gliomas & PDX susceptibility to TMZ**.

**(A)** Pictures show immunofluorescence of NSUN6 across GBM (MES83, GBM39, GBM44, GBM6) with NSUN6 (green) and nuclear staining DAPI (blue). **(B)** Pictures show immunofluorescence of 5mC (green) and nuclear staining DAPI (blue) across GBM (MES83, GBM39, GBM44, GBM6).

**Supplementary Figure 2: TATA binding proteins and RNAPOLII accumulates in NSUN6 edited cells**.

**(A)** Graph shows one phase decay half-life curve comparing TATA binding protein (TBP) mRNA stability in SgNT (black) and SgNSUN6 (yellow) U251 total RNA. **(B)** Graph shows one phase decay half-life curve comparing RNA POLII (POLR2A) mRNA stability in SgNT (black) and SgNSUN6 (green) U251 total RNA. **(C)** Dot blot shows total RNA treated with 5 aza deoxy cytidine and stained with methylene blue and anti-5mC. **(D)** Bar charts represents fold change of GTF (general transcription factors) enrichment normalized against GAPDH of Azacytidine treated cells total RNA.

**Supplementary Figure 3: Multivariate analysis shows no correlation between MGMT and NSUN6 expression**.

**(A&B)** Chart shows multivariate analysis comparing the expression of NSUN6 against MGMT for patients treated with temozolomide (Statistical test=Pearsons /spearman’s)

---

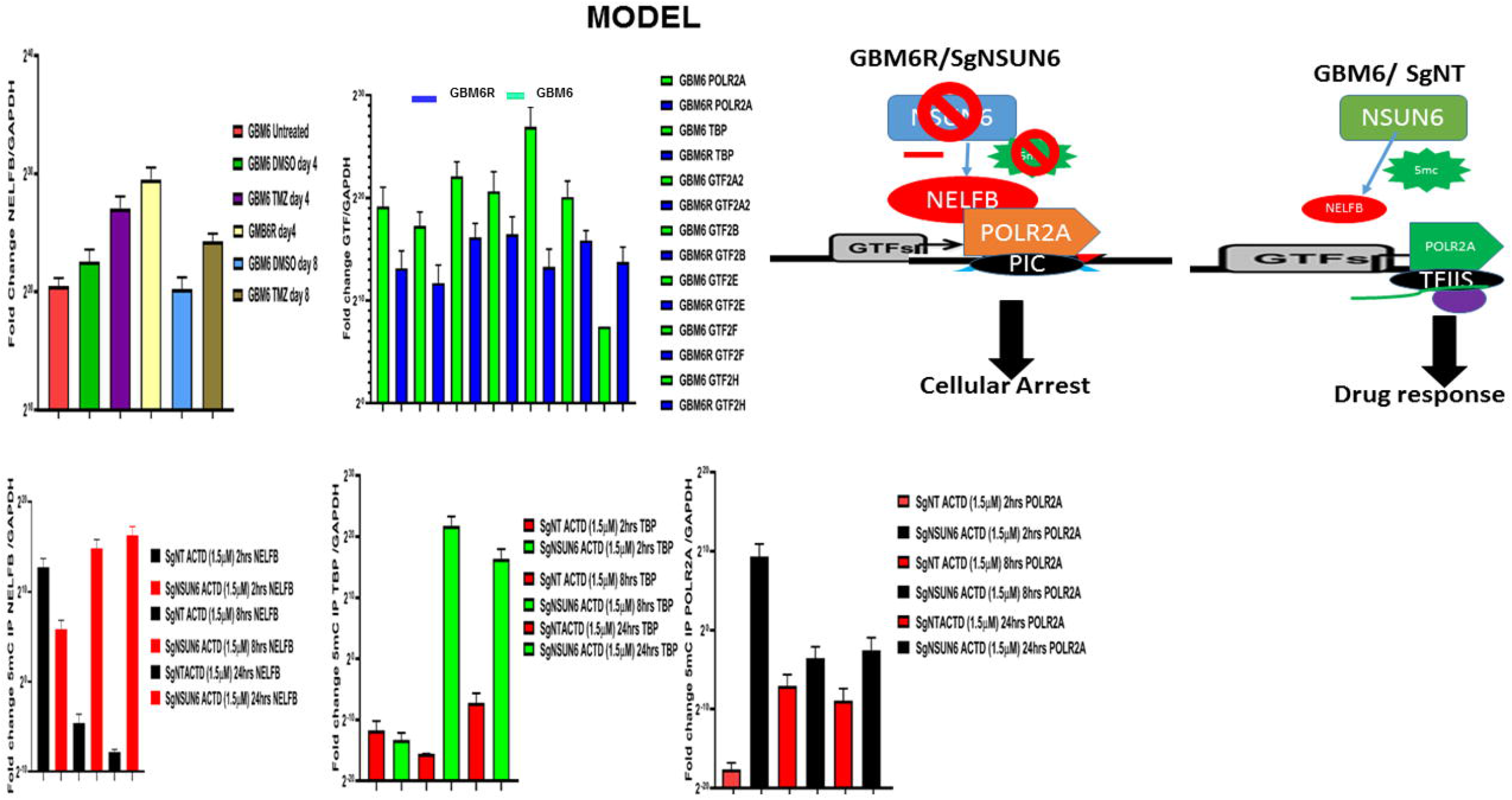

## Notes

https://zenodo.org/record/5167143#.YQ6tHYhKiM8

### Competing Interest Statement

The authors have declared no competing interest.

