## Supplementary Figure 1 for "NSUN6, an RNA methyltransferase of 5-mC controls glioblastoma response to Temozolomide (TMZ) via NELFB and RPS6KB2 interaction"

Supplementary Fig1:Expression of NSUN6 correlates with it's RNA methylation mark 5mC in the cytoplasm of gliomas & PDX susceptibility toTMZ

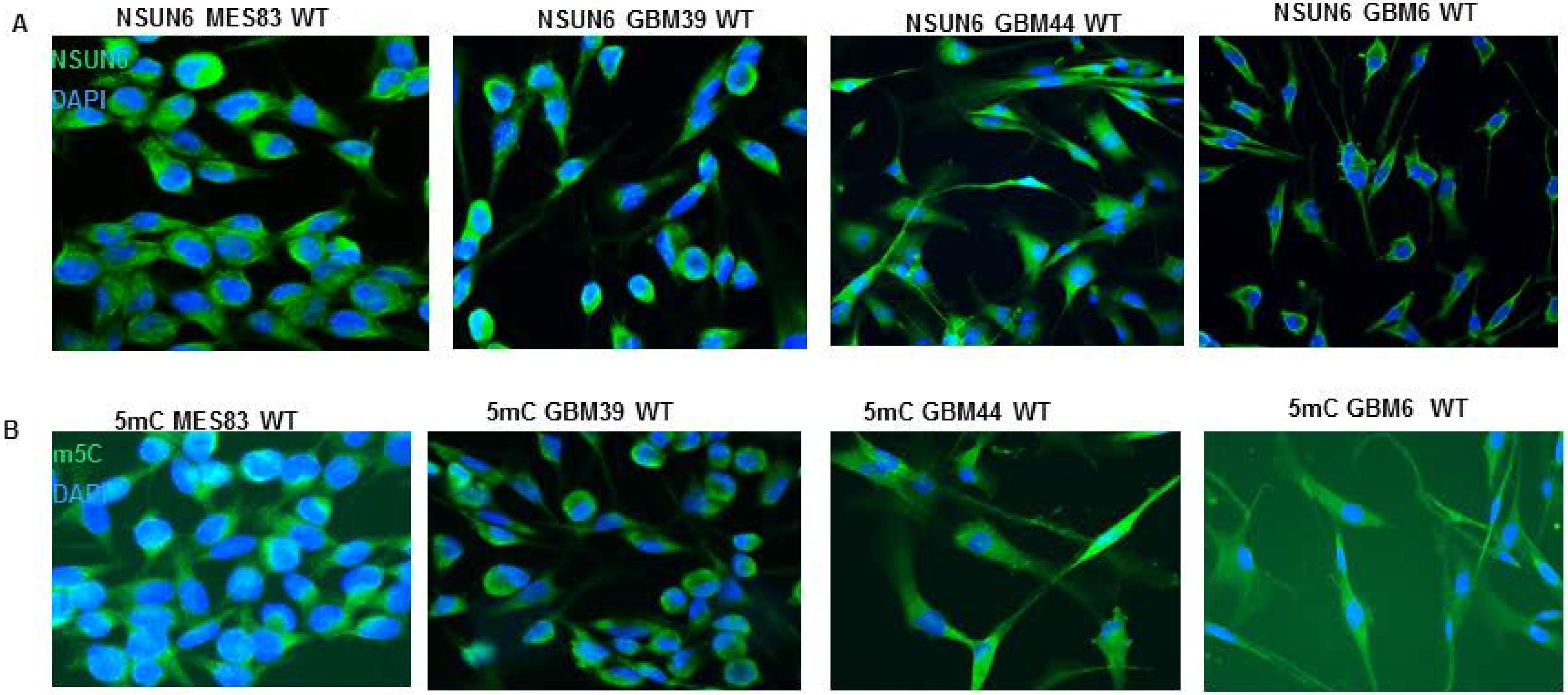
