## Supplementary figures and images for "NSUN6, an RNA methyltransferase of 5-mC controls glioblastoma response to Temozolomide (TMZ) via NELFB and RPS6KB2 interaction"

### Supplemntary Figure 2

Supplementary Fig 2: TATA binding proteins and RNAPOLII accumulates in NSUN6 edited cells

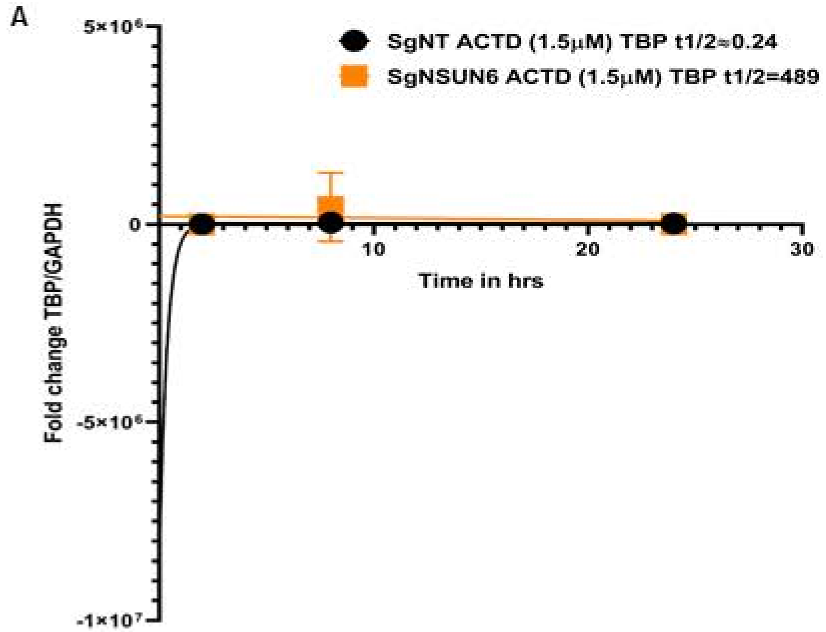


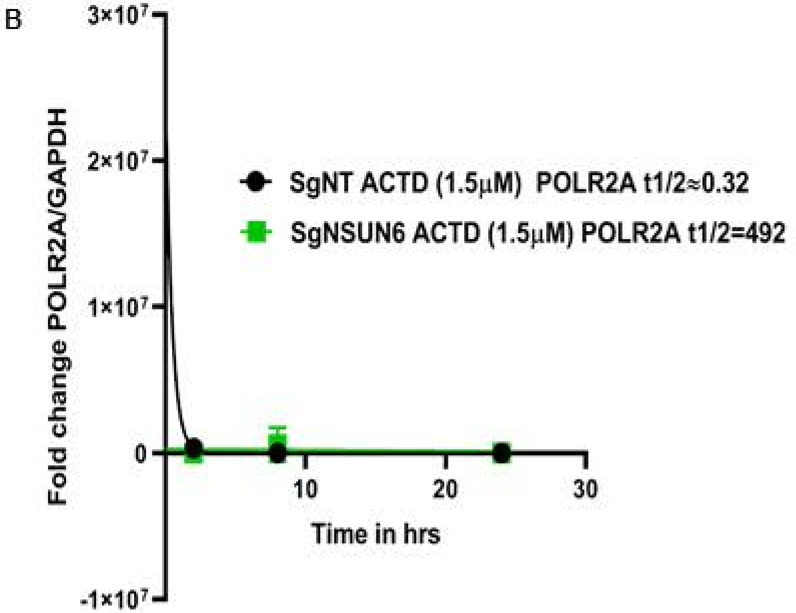


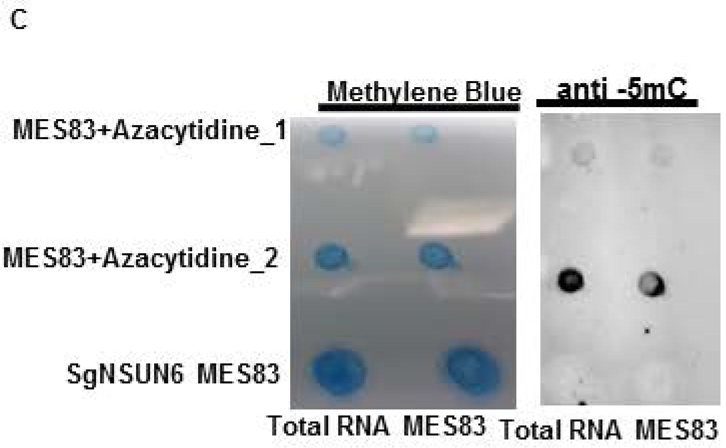


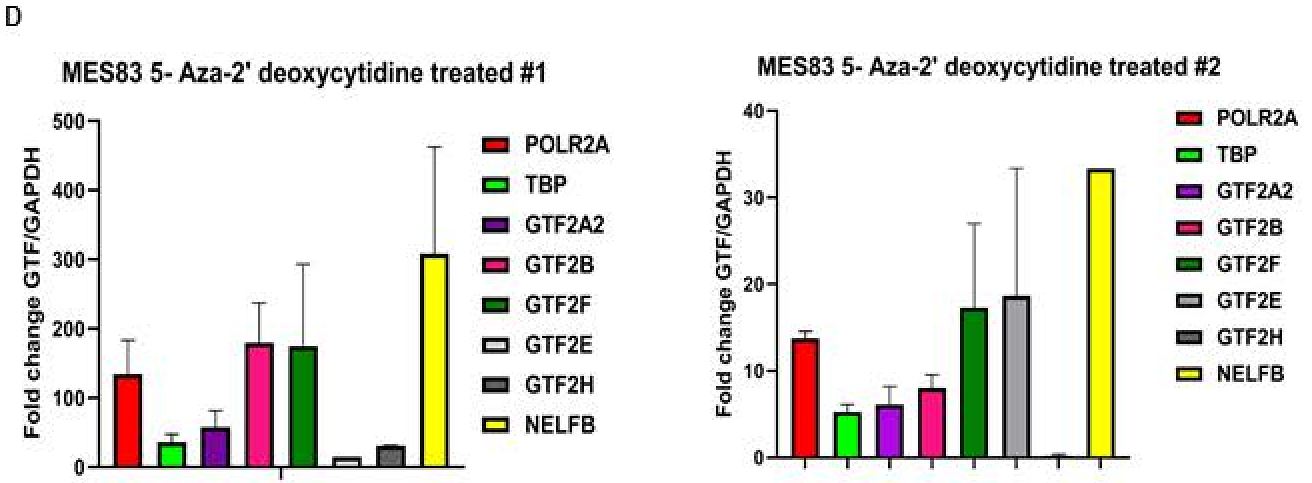
