## Supplementary Figure 3 for "NSUN6, an RNA methyltransferase of 5-mC controls glioblastoma response to Temozolomide (TMZ) via NELFB and RPS6KB2 interaction"

**Supplementary Fig 3: Multivariate analysis shows no correlation between MGMT and NSUN6 expression.**

A **Pearson's correlation**

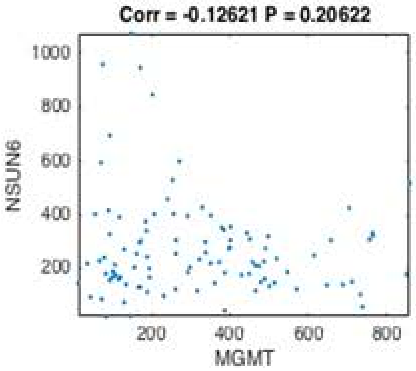


B **Spearman's correlation**

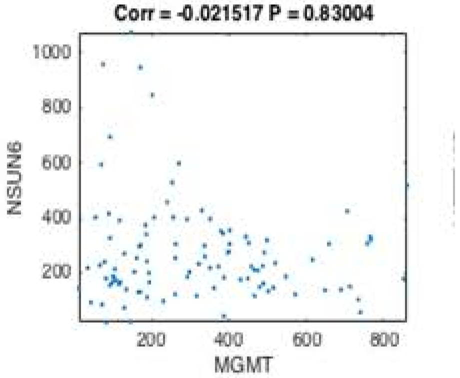
